# Elevated SUN1 promotes aging-related polarity defects through mechanical coupling microtubules to the nuclear lamina

**DOI:** 10.1101/2025.01.15.633301

**Authors:** Yutao Li, Mengqi Chen, Susumu Antoku, Junjun Ding, Kaiyao Huang, Gregg G. Gundersen, Wakam Chang

## Abstract

In migratory fibroblasts, the front-rear polarity required for cell migration is defined by an anterior centrosome relative to the nucleus. To achieve this polarity, actin cables drive the nucleus backward by coupling to nuclear membrane proteins nesprin-2G and SUN2. Aging disrupts this cell polarity by increasing the protein levels of SUN1, a SUN2 homolog. Here, we investigated the molecular mechanisms behind this disruption and found that the dominant negative effect of SUN1 and progerin, an aging-related lamin A variant, required direct SUN1-lamin A interaction. Microtubule interaction and force transmission through a nesprin, identified as nesprin-2, are crucial for SUN1’s effect. We further discovered that stable microtubules are both necessary and sufficient to inhibit cell polarity. Using SUN1-SUN2 chimeric proteins, we demonstrated that the SUN domains determine their roles in cell polarization. Our findings reveal how elevated SUN1 disrupts cell polarity through coupling microtubule and nuclear lamina, emphasizing the impact of altered microtubule stability and nuclear mechanotransduction in aging.

## Introduction

Aging is a fundamental biological phenomenon resulting from the accumulation of deficits in an organism. While these deficits are most obvious at the organ and tissue levels, at their core, they are rooted in molecular and cellular alterations. Cells are the basic units of life, and the elucidation of aging-related cellular defects has contributed tremendously to our current understanding of the mechanisms of aging [1]. In addition to “canonical” defects that are widely studied, like DNA damage and metabolic alterations, more and more cellular processes have been revealed to contribute to aging [1,2]. Cell polarity, the asymmetric distribution of molecules and structures, is a universal feature of cells. The association between cell polarity and aging has been characterized in budding yeast and, more recently, stem cells [3,4]. In both systems, abnormal polarity contributes to aging by affecting differential inheritance and asymmetric cell division. However, cell polarity is critical to all aspects of cells, and its deficiencies may give rise to other aging defects, like impaired tissue architecture and cell migration [5,6].

Impaired cell migration is a long-established aging phenotype and contributes to the decline of wound healing during aging [7]. Cell polarity establishes the direction of movement in migratory cells and is in part determined by the centrosomal position relative to the nucleus. In fibroblasts, cell polarity is induced by lysophosphatidic acid (LPA), which binds to its receptor on the plasma membrane to activate the small GTPase Cdc42. Cdc42, a master regulator of cell polarity, activates an MRCK-myosin-actin pathway to move the nucleus backward and a PKCξ-Par6-Par3-dynein-microtubule pathway to center the centrosome in the cell [8,9]. Disruptions in these pathways lead to a centered nucleus and/or a misoriented centrosome, collectively referred to as polarity defects, resulting in migration defects. Many of the proteins in these pathways, including the LPA receptor, Cdc42, and Par3, have been implicated in aging [10–12].

Rearward nuclear movement requires force transmission from actin filaments to the nucleus, which is mediated by linker of nucleoskeleton and cytoskeleton (LINC) complexes composed of nesprin on the outer nuclear membrane (ONM) and Sad-1 and UNC-84 (SUN) domain proteins on the inner nuclear membrane (INM) [13,14]. The LINC complexes connect the cytoskeleton and the nuclear lamina through three interactions separated by the nuclear membranes. Moving from the cytoplasm to the nucleus, the first of these interactions is between a nesprin and one or more cytoskeletal elements. Specificity for actin filaments (nesprin-1/2), microtubules (nesprin-1/2/4), and intermediate filaments (nesprin-3) is determined by motifs in the specific nesprin and associating proteins [13]. The next is the interaction between the nesprin KASH (Klarsicht, ANC-1, and Syne homology) domain and the SUN domains of SUN proteins in the lumen of the ONM and INM [15,16]. The last is the interaction between the nucleoplasmic domains of SUN proteins and lamins of the nuclear lamina. Somatic cells express four nesprins (nesprin-1 to −4) and two SUN proteins (SUN1 and SUN2). Among them, nesprin-2G, a giant isoform of nesprin-2, and SUN2 are involved in fibroblast nuclear movement by connecting actin filaments to lamin A [17–20].

The nesprin-2G and SUN2-dependent movement of the nucleus is inhibited when SUN1 expression is experimentally elevated [21]. This “transdominant inhibition” presumably occurs because both SUN proteins bind to nesprin-2G, but SUN1 promotes nesprn-2G’s interaction with microtubules rather than actin filaments [21]. Elevated SUN1 levels are observed in several conditions, including Hutchison Gilford progeria syndrome (HGPS) and physiological aged human fibroblasts, correlating with impaired migratory polarity in these cells [22,23]. HGPS is caused by the expression of progerin, a farnesylated mutant of lamin A [24,25]. Ectopic expression of progerin is sufficient to inhibit nuclear movement by increasing SUN1 levels [23]. SUN1 is also elevated in a mouse model of HGPS and in stem cells of normally aged mice [22,26]. Notably, reducing SUN1 expression improves phenotypes in all these cases. In human fibroblasts, nuclear movement and resulting polarity are restored by SUN1 knockdown [23]. In mice, SUN1 knockout significantly improves aged phenotypes, cell stemness, and lifespan [22,26].

The mechanism by which elevated SUN1 causes aging-related phenotypes remains unclear. It is also perplexing how the nuclear SUN proteins regulate cytoplasmic nucleus-cytoskeletal interactions. In this study, we use the relatively simple system of cell polarity acquisition in wounded monolayers of fibroblasts and investigate SUN1’s direct molecular interactions. We demonstrate that the SUN1-lamin A interaction is required for both SUN1 and progerin to impede cell polarization. The SUN1-nesprin interaction and its ability to transmit strong forces are also essential for both SUN1 and stable microtubules to hinder cell polarity, suggesting the involvement of microtubules and mechanotransduction in this process. By employing SUN1-SUN2 chimeras, we showed that the cell polarity-promoting activity of SUN2 and the inhibiting activity of SUN1 are determined by their SUN domains rather than their coiled-coil or nucleoplasmic domains. Altogether, our results establish a new paradigm for how LINC complex specificity for different cytoskeletal elements is established and show how changes in SUN1 or progerin levels contribute to age-dependent polarity defects.

## Results

### Progerin inhibits cell polarity through direct interaction with SUN1

In a previous study, we found that depletion of SUN1 reversed polarity defects in fibroblasts from patients with HGPS [23]. These primary cells accumulate numerous cellular defects [27], making them less than ideal to study the mechanistic roles of SUN1. Ectopic expression of progerin in wildtype fibroblasts is sufficient to interfere with cell polarization [23], providing an opportunity to test how SUN1 contributes to progerin-induced defects. In NIH3T3 fibroblasts stably expressing myc-tagged progerin, we used shRNAs to knock down SUN1. Depletion of SUN1 restored nuclear movement and centrosome orientation in these cells (Fig. S1), confirming that SUN1 is required for polarity defects caused by progerin.

Progerin alters a myriad of cellular pathways via largely unknown molecular interactions [27,28]. SUN1 directly interacts with lamin A and binds to both prelamin A and progerin with increased affinities [15,29]. However, whether SUN1 mediates the effects of progerin through their direct interaction is unknown. To investigate if the interaction is required for the SUN1-dependent inhibition of cell polarity, we introduced an R527P point mutation into progerin, which was previously shown to reduce interaction between SUN1 and mature lamin A [16]. The mutant protein was expressed at a similar level as progerin (Fig. 1A). We confirmed that the R527P mutation reduced the association between progerin and SUN1 (Fig. 1B). We then examined their localization and effect on cell polarity. Myc-progerin^R527P^ and myc-progerin both localized to the nuclear envelope as expected (Fig. S2A). However, unlike myc-progerin, myc-progerin^R527P^ did not interfere with cell polarity (Fig. 1C-E). Reduction in cell migration by progerin expression, as measured by wound healing experiments, was also reversed by the mutation (Fig. S2B, C). These results demonstrated that progerin induces cell polarity and cell migration defects through its interaction with SUN1.

**Figure 1.**
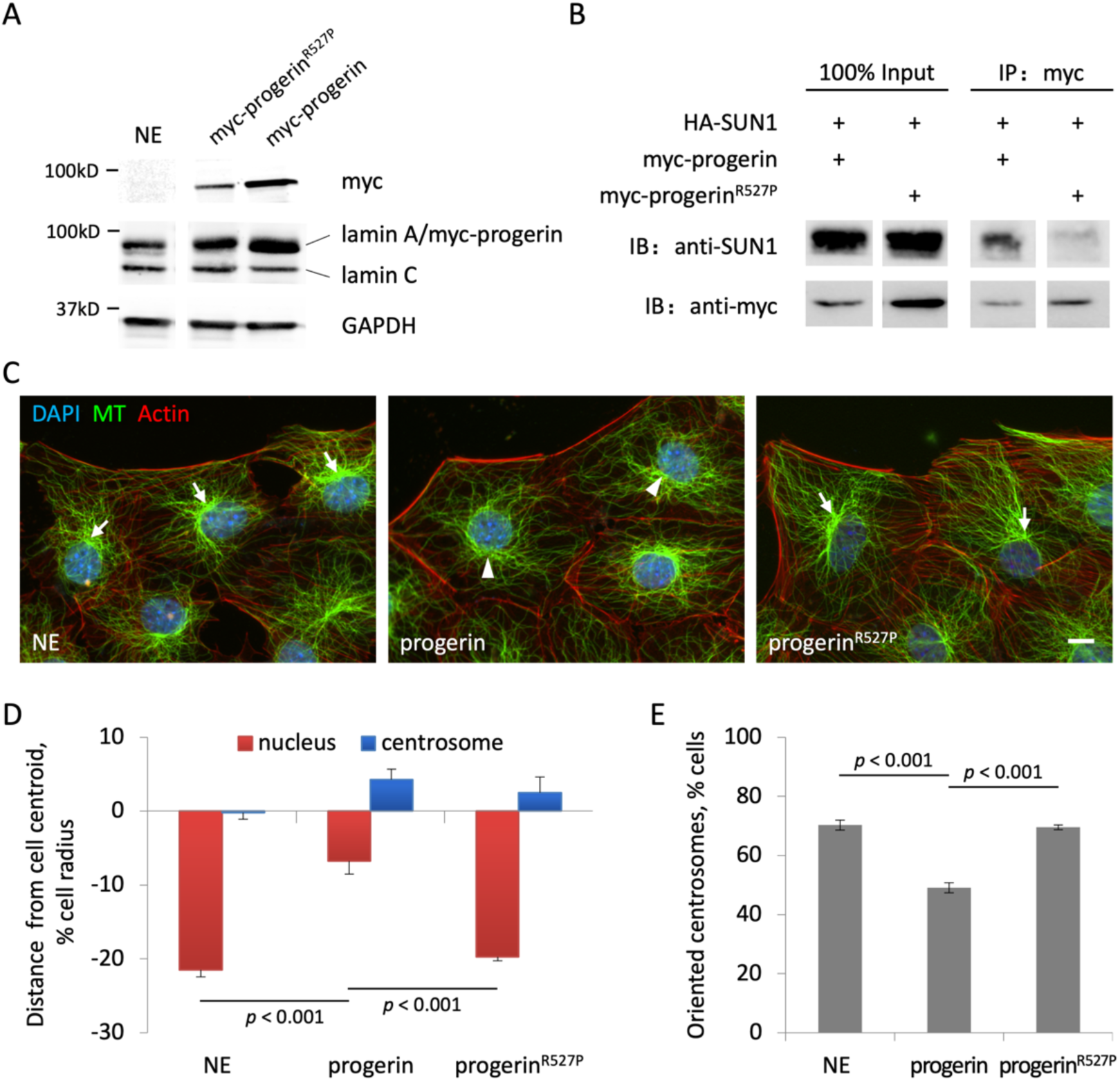
Progerin must interact with SUN1 to inhibit rearward nuclear movement and cell polarization. **A)** Western blot showing the expression of myc-progerin and myc-progerin^R527P^ in NIH3T3 fibroblasts. **B)** Western blot of input and pellet of anti-myc immunoprecipitation using lysates from cells expressing HA-SUN1 and myc-progerin/progerin^R527P^. **C)** Representative images of wounded monolayers of LPA-induced non-expressing NIH3T3 fibroblasts (NE) and cells expressing myc-progerin and myc-progerin^R527P^. Cells were fixed 2 h after 10 µM LPA was added and stained for α-tubulin (MT), phalloidin (Actin), and DNA (DAPI). Arrows, centrosomes oriented towards the wound. Arrowheads, misoriented centrosomes. Bar, 10 µm. **D)** Nuclear and centrosome positions along the front-back axis for cells treated as in panel C. The numbers represented the distances from the cell centroid, adjusted relative to the cell radii. Positive values: toward the wound edge, negative values: away from the wound edge. **E)** Levels of oriented centrosomes for cells treated as in panel C. Centrosome orientation by chance is 33%. All values are mean ± SEM. Experiments were repeated three-times (*N* = 3) and at least 90 cells (*n* ≥ 90) were analyzed in each condition. *p*-values are by one-way ANOVA with a post-hoc Tukey test.

### SUN1’s interaction with lamin A is required for its inhibition of cell polarity

We investigated whether SUN1’s inhibitory effects on cell polarity require its interaction with lamin A. We deleted the first 138 amino acids of SUN1 (Fig. S3A), which are responsible for its binding to lamin A, but not to emerin and MAJIN [30,31]. We tagged the resultant fragment (SUN1ΔN) with myc and expressed it in NIH3T3 fibroblasts. Myc-SUN1ΔN localized normally to the nuclear envelope (Fig. S3B), confirming that this domain is not required for the localization of SUN1 [16]. Despite myc-SUN1ΔN being expressed at higher levels than myc-SUN1 (Fig. S3A), overall cell polarity, as indicated by both rearward nuclear movement and centrosome orientation, was significantly improved compared to cells expressing myc-SUN1 (Fig. 2A-C). These results demonstrate that the interaction with lamin A/C is critical for overexpressed SUN1 to inhibit cell polarity.

**Figure 2.**
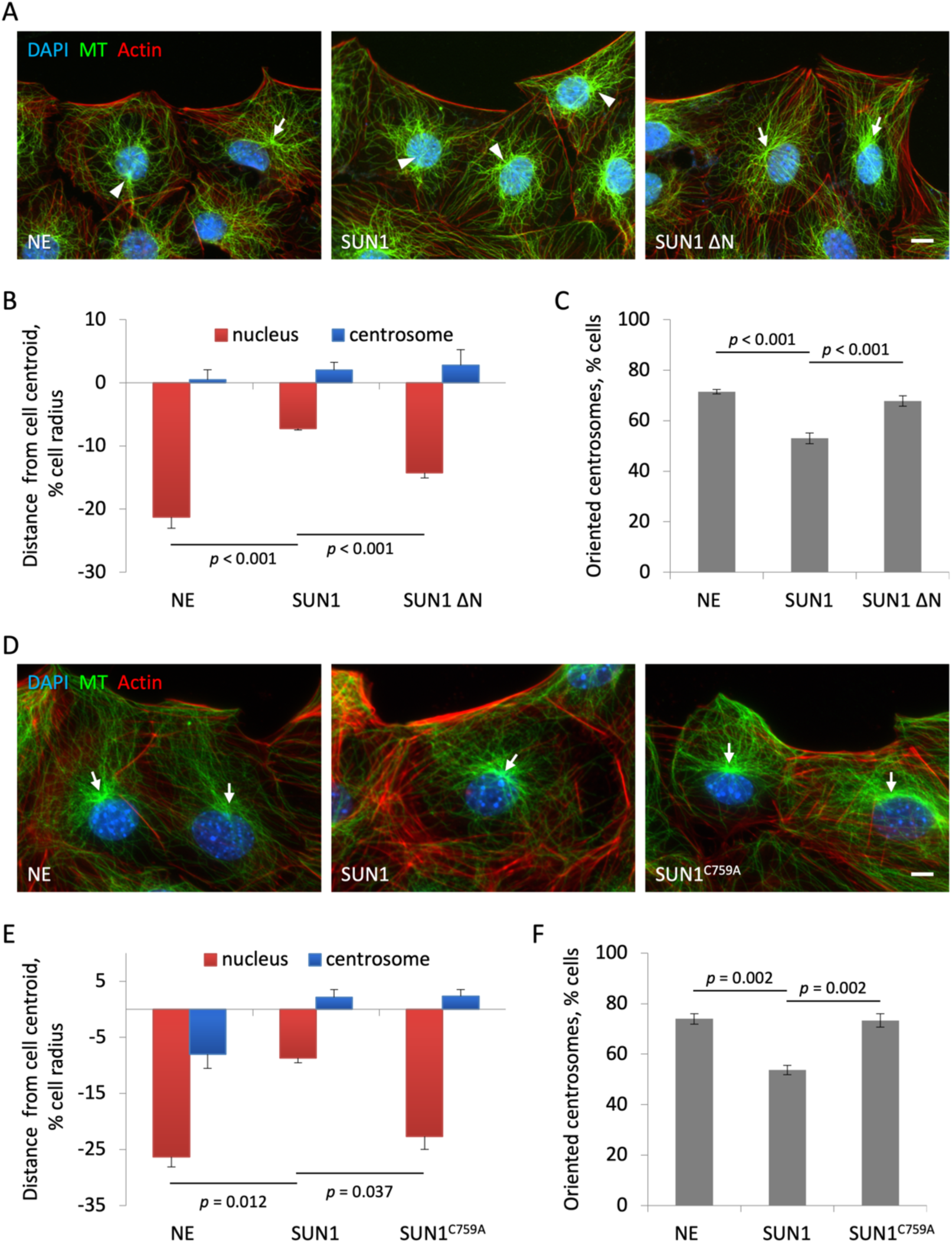
SUN1’s lamin A-interacting domain and covalent interaction with nesprins are necessary for the inhibition of rearward nuclear movement. **A)** Representative images of wounded monolayers of LPA-stimulated non-expressing NIH3T3 fibroblasts (NE) and fibroblasts expressing myc-SUN1 and myc-SUN1ΔN. **B, C)** Nuclear and centrosome positions (B) and levels of oriented centrosomes (C) for cells as in panel A. **D)** Representative images of LPA-stimulated non-expressing NIH3T3 fibroblasts and fibroblasts expressing myc-SUN1 and myc-SUN1^C759A^. **E, F)** Nuclear and centrosome positions (E) and levels of oriented centrosomes (F) for cells as in panel D. Bars, 10 µm. *p*-values are by one-way ANOVA with a post-hoc Tukey test (*N* = 3, *n* ≥ 90).

### Force transmission through nesprins-2G is required for SUN1 to inhibit cell polarity

SUN1 binds to nesprins in the ONM to form the LINC complex, and most SUN1 functions rely on nesprins. We thus asked if the binding between SUN1 and nesprins is required for SUN1’s inhibition of cell polarity. The SUN-KASH interaction involves the insertion of the KASH peptide into a groove between two SUN domains and the formation of a disulfide bond between the KASH and SUN domains [32–34]. Molecular dynamics simulations and nuclear positioning assays suggest that the disulfide bond is critical for the transmission of strong forces to move the nucleus [35,36]. We overexpressed myc-tagged SUN1 carrying a C759A mutation in the SUN domain that prevents the formation of the disulfide bond [35]. SUN1^C759A^ localized correctly to the nuclear envelope (Fig. S4A) but exerted no inhibitory effect on cell polarity (Fig. 2D-F). Therefore, the covalent interaction with nesprins and the ability to transmit strong forces are essential for SUN1 to disrupt cell polarity. Wound healing experiments showed that cell migration is slightly, but significantly, inhibited by overexpressed SUN1 (Fig. S4B, C). Consistent with the lack of polarity defects, cell migration was not affected in cells expressing SUN1^C759A^.

SUN1 and SUN2 are capable of binding the same nesprins [37,38], and there is evidence that they compete for nesprins [21,23,39]. Thus, elevated SUN1 may interfere with SUN2-nesprin-2G function by competing for nesprin-2G. To test if reduced SUN2-nesprin-2G-actin interaction is the major contributing factor to impaired nuclear movement caused by elevated SUN1, we expressed mini-nesprin-2G, which rescues nuclear movement in nesprin-2G-depleted fibroblasts [17]. Expression of mini-nesprin-2G in SUN1-overexpressing cells improved rearward nuclear movement (Fig. 3A-C), indicating that boosting the nesprin connection to the actin cytoskeleton enhances cell polarity. However, the improvement was only partial, suggesting that SUN1 inhibits nuclear movement through a mechanism other than competing with SUN2.

**Figure 3.**
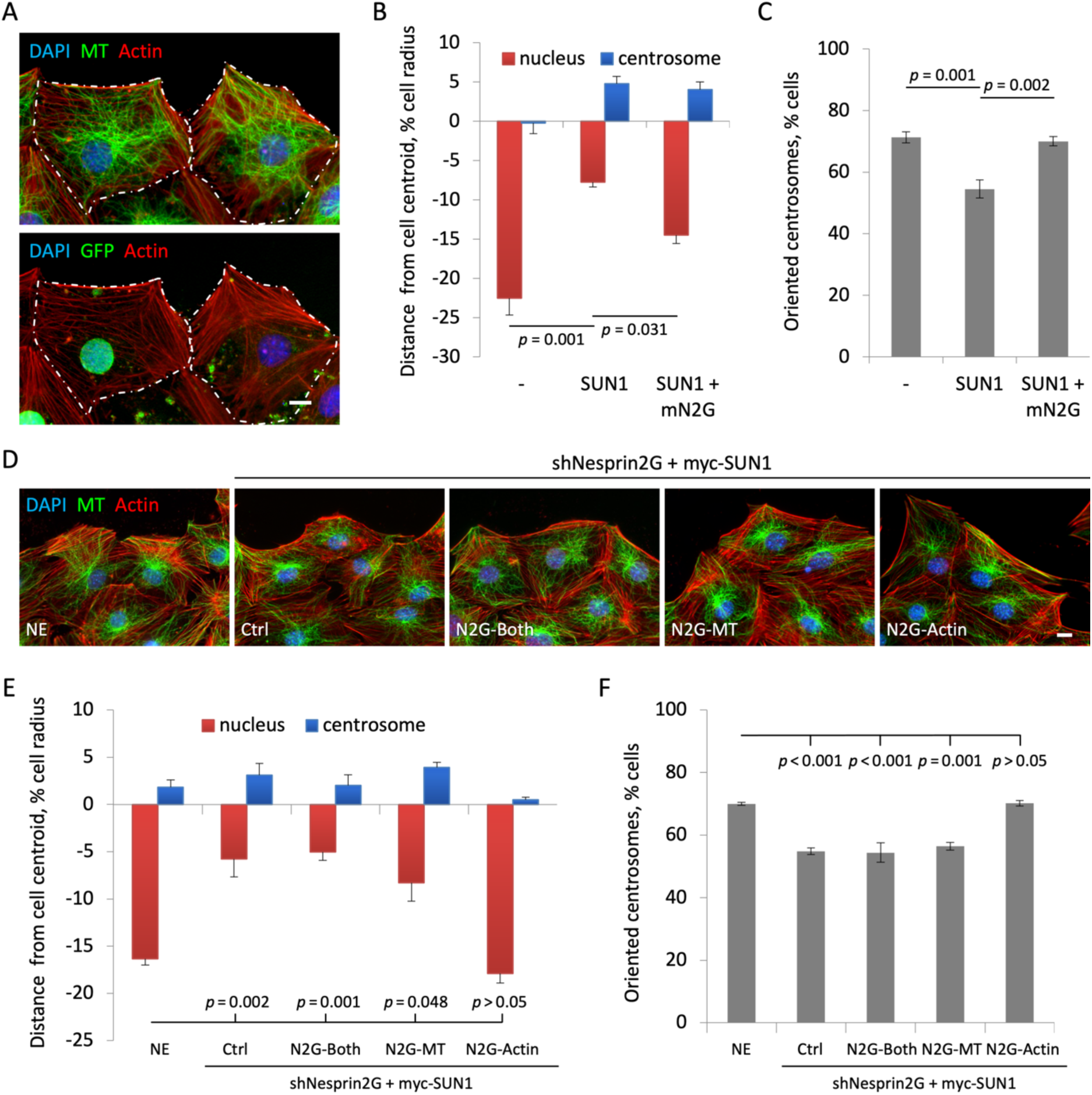
Enhancing nuclear-actin association with mini-nesprin-2G expression only partially rescues rearward nuclear movement. **A)** Images of wounded monolayers of LPA-stimulated NIH3T3 fibroblasts expressing both myc-SUN1 and GFP-mN2G (left) or myc-SUN1 alone (right). **B, C)** Nuclear and centrosome positions (B) and levels of oriented centrosomes (C) for cells expressing the indicated proteins. **D)** Images of LPA-stimulated non-expressing cells (NE), cells expressing shNesprin-2G and myc-SUN1 (Ctrl), and cells additionally expressing the indicated GFP-tagged nesprin-2G fragments. **E, F)** Nuclear and centrosome positions (E) and levels of oriented centrosomes (F) for cells in panel D. Bar, 10 µm. *p*-values are by one-way ANOVA with a post-hoc Tukey test (*N* = 3, *n* ≥ 90).

We then asked if nesprin-2G and nesprin-2G-mediated SUN1-microtubule interaction are involved in polarity inhibition by SUN1. We used shRNA to deplete nesprin-2G in SUN1-overexpressing cells (Fig. S5A, B) and expressed GFP-tagged nesprin-2G fragments that bind to either actin filaments (N2G-Actin, same as mini-nesprin-2G), microtubules (N2G-MT), or both (N2G-Both) [21]. All these fragments localized to the nuclear envelope as expected (Fig. S5C, D). Both N2G-Actin and N2G-Both were shown to support actin-based nuclear movement in the absence of nesprin-2G [21]. However, only N2G-Actin, and not N2G-MT or N2G-Both, rescued nuclear movement in nesprin-2G-depleted, SUN1-overexpressing cells (Fig. 3D-F). The differing effects between N2G-Actin and N2G-Both strongly suggest that the microtubule-interacting region of nesprin-2G is required for SUN1 to inhibit nuclear movement.

### SUN1 inhibits cell polarity through microtubules

When bound to SUN1, nesprin-2G engages microtubules through dynein and kinesin-1 motors [40,21]. Treatment with HPI-4, a dynein inhibitor, rescues cell polarity in both physiologically and pathologically aged human fibroblasts, leading us to propose that the upregulated SUN1 levels in aging increase the association between microtubules and the nucleus [23]. Supporting this hypothesis, rearward nuclear movement and centrosome orientation are restored in SUN1-expressing cells pretreated with HPI-4 for 1 h (Fig. 4A-C).

**Figure 4.**
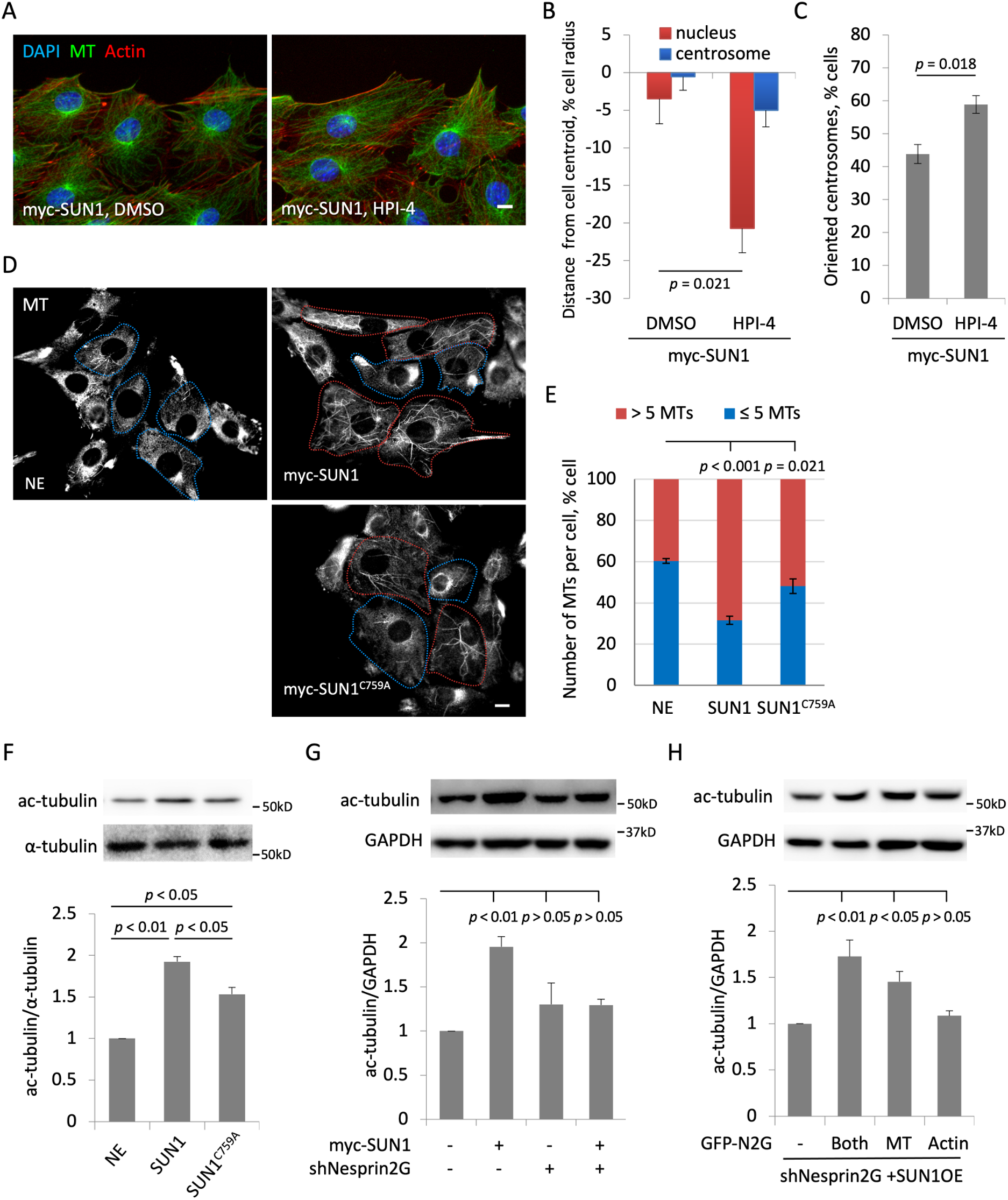
SUN1 inhibits rearward nuclear movement through microtubules and enhances microtubule stability through the microtubule-interaction region nesprin-2. **A)** Representative images of wounded monolayers of LPA-stimulated myc-SUN1 expressing NIH3T3 fibroblasts pretreated with DMSO or 28 µM HPI for 1 h. **B, C)** Nuclear and centrosome positions (B) and levels of oriented centrosomes (C) for cells as in panel A. **D)** Representative images of α-tubulin staining (MT) in non-expressing cells (NE) and NIH3T3 fibroblasts expressing the indicated proteins after 15 min of nocodazole treatment (3.3 μM). **E)** Number of remaining microtubules for cells in panel D. **F)** Western blot (top) and quantification (bottom) of acetylated (ac-) and total α-tubulin in wildtype cells and cells expressing the indicated myc-tagged proteins. **G)** Expression of acetylated α-tubulin in NIH3T3 fibroblasts expressing myc-SUN1 and/or shNesprin2G as indicated. **H)** Expression of acetylated α-tubulin in NIH3T3 fibroblasts expressing myc-SUN1, shNesprin-2G, and the indicated nesprin-2G fragment. Bars, 10 µm. *p*-values are by Student’s t-test (B, C, *N* = 3, *n* ≥ 90), Chi-Square test (E, *N* = 3), and one-way ANOVA with a post-hoc Tukey (F, *N* = 3) or Dunnett test (G, H, *N* = 3).

Fibroblasts from HGPS patients have elevated levels of stable microtubules around the nucleus, indicating a role for SUN1 engaging microtubules with the nucleus [23]. Consistent with this, a recent study showed that SUN1 contributes to microtubule stability encircling nuclei in endothelial cells [41]. The tremendous numbers of microtubules in NIH3T3 fibroblasts hindered our detection of changes in microtubule numbers, so we used brief treatment with nocodazole to remove dynamic microtubules. Compared to wildtype cells, SUN1-overexpresing cells retained significantly more microtubules after nocodazole treatment (Fig. 4D, E). Unexpectantly, SUN1^C759A^, which cannot form a covalent LINC complex with nesprins, also stabilized microtubules (Fig. 4D, E). A second marker of stable microtubules, α-tubulin acetylation, was increased by both SUN1 and SUN1^C759A^ (Fig. 4F).

To test whether the increased microtubule stability induced by SUN1 expression depended on its LINC complex function, we depleted nesprin-2G. SUN1-induced α-tubulin acetylation was significantly reduced in nesprin-2G-depleted cells (Fig. 4G). This reduction could be reversed by N2G-MT and N2G-Both, which contain the microtubule-binding regions, but not by N2G-Actin (Fig. 4H). Thus, SUN1 enhances microtubule stability through nesprin-2G, and this capability may not depend solely on its ability to form a covalent LINC complex.

To test if microtubule stability is indeed involved in polarity inhibition, we used remodelin, a chemical inhibitor of N-acetyltransferase 10 (NAT10) shown to reduce microtubules and rescue abnormal nuclear morphology in cells from HGPS patients [42]. Remodelin treatment reduced α-tubulin acetylation (Fig. 5A, S6A, B) and significantly rescued rearward nuclear movement in SUN1-overexpressing cells (Fig. 5B, C, S6C), suggesting that SUN1 inhibits cell polarity through stable microtubules. Next, we asked if stabilizing microtubules is sufficient to impair cell polarization. We pretreated wildtype NIH3T3 fibroblasts with a low dosage (2 nM) of the microtubule-stabilizing drug Paclitaxol (Taxol) for 30 min before triggering nuclear movement. This Taxol treatment inhibited nuclear movement and centrosome orientation (Fig. 5D-F). We then investigated whether the inhibitory effect of stabilizing microtubules on cell polarity requires SUN1. Using two specific shRNAs against SUN1 to reduce SUN1 expression (Fig. S6D), we found that nuclear movement was significantly improved (Fig. 5D-F). Cotreating cells with HPI-4 also counteracted the polarity inhibition by Taxol (Fig. S7). Therefore, stable microtubules alone are not sufficient; their association with the nucleus is also necessary to inhibit nuclear movement. Combined, these results show that elevated SUN1 and stable microtubules function together to inhibit cell polarity.

**Figure 5.**
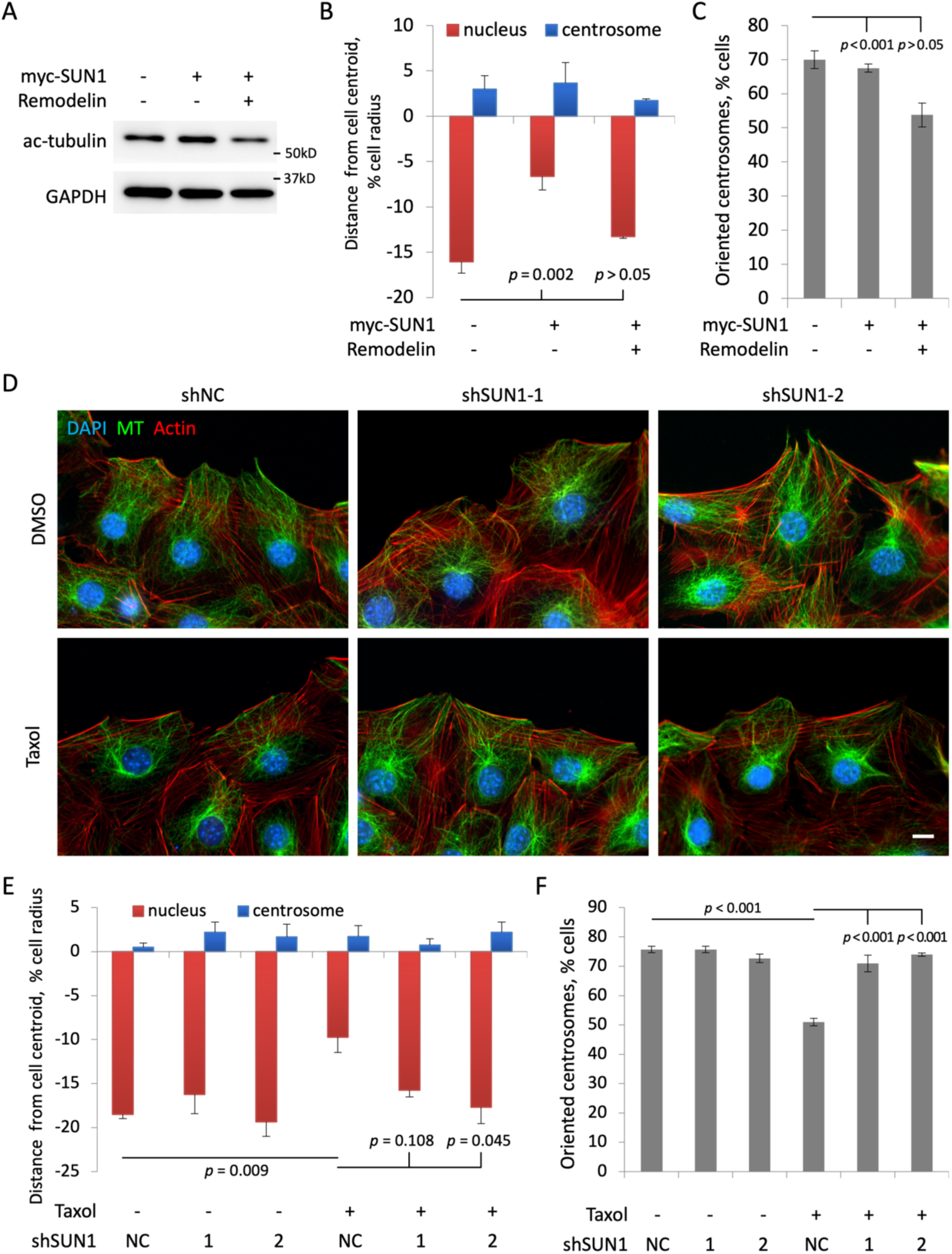
Stable microtubules inhibit actin-dependent nuclear movement in a SUN1-dependent manner. **A)** Acetylated α-tubulin (ac-tubulin) levels in NIH3T3 fibroblasts with and without expression of myc-SUN1. Cells were treated with either DMSO or remodelin (10 µM) for 24 h. GAPDH serves as a loading control. **B, C)** Nuclear and centrosome positions (B) and levels of oriented centrosomes (C) for LPA-stimulated cells treated as indicated. **D)** Representative images of LPA-stimulated NIH3T3 fibroblasts expressing the non-coding (NC) or SUN1-specific shRNAs and pretreated with DMSO or 2nM Taxol for 30 min. Bar, 10 µm. **E, F)** Nuclear and centrosome positions (E) and levels of oriented centrosomes (F) for cells treated as in panel D. *p*-values are by one-way ANOVA with a post-hoc Tukey test (*N* = 3, *n* ≥ 90).

### SUN domains determine the functions of SUN1 and SUN2 in cell polarity

Somatic cells express two SUN proteins, SUN1 and SUN2. Both interact with nesprin-2G but have distinct functions during fibroblast nuclear movement [38,21]. SUN2 and nesprin-2 are required for actin-dependent nuclear movement, while SUN1-nesprin-2G mediates microtubule-dependent nuclear. How SUN interaction specifies nesprin-2’s interaction with either actin filaments or microtubules is unknown.

We took advantage of the MT-dependent, polarity-inhibiting effect of SUN1 overexpression to test whether functional specificity resided in one of the three domains of the SUN protein: its amino-terminal nucleoplasmic domain (NP), which binds lamins and is the most diverse between SUN1 and SUN2; its middle coiled-coil (CC) region, which mediates trimerization; or its C-terminal SUN domain, which binds nesprins (Fig. 6A). We created chimeras in which the nucleoplasmic domains of SUN proteins were swapped (Fig. 6B). After retroviral infection, these chimeras were expressed in virtually all cells and, by immunofluorescence, both localized to the nuclear envelope as expected and were expressed at equivalent levels (Fig. S8A). Expression of a SUN1 chimera with the nucleoplasmic domain of SUN2 (S1-CC1-NP2) blocked rearward nuclear movement and centrosome orientation, whereas expression of a SUN2 chimera with SUN1’s nucleoplasmic domain (S2-CC2-NP1) did not (Fig. 6C-E). These results suggest that the nucleoplasmic domains of SUN protein do not have critical roles in determining nesprin-2G’s functional specificity toward actin or microtubules. Next, we created chimeras in which the SUN domains were swapped (Fig. 6B). These chimeras also expressed equivalently and localized correctly (Fig. S8A). The chimera with the SUN1 domain (S1-CC2-NP2), but not the one with the SUN2 domain (S2-CC1-NP), inhibited nuclear movement and centrosome orientation (Fig. 6C-E), indicating that the SUN domain plays a determinant role in functional specificity.

**Figure 6.**
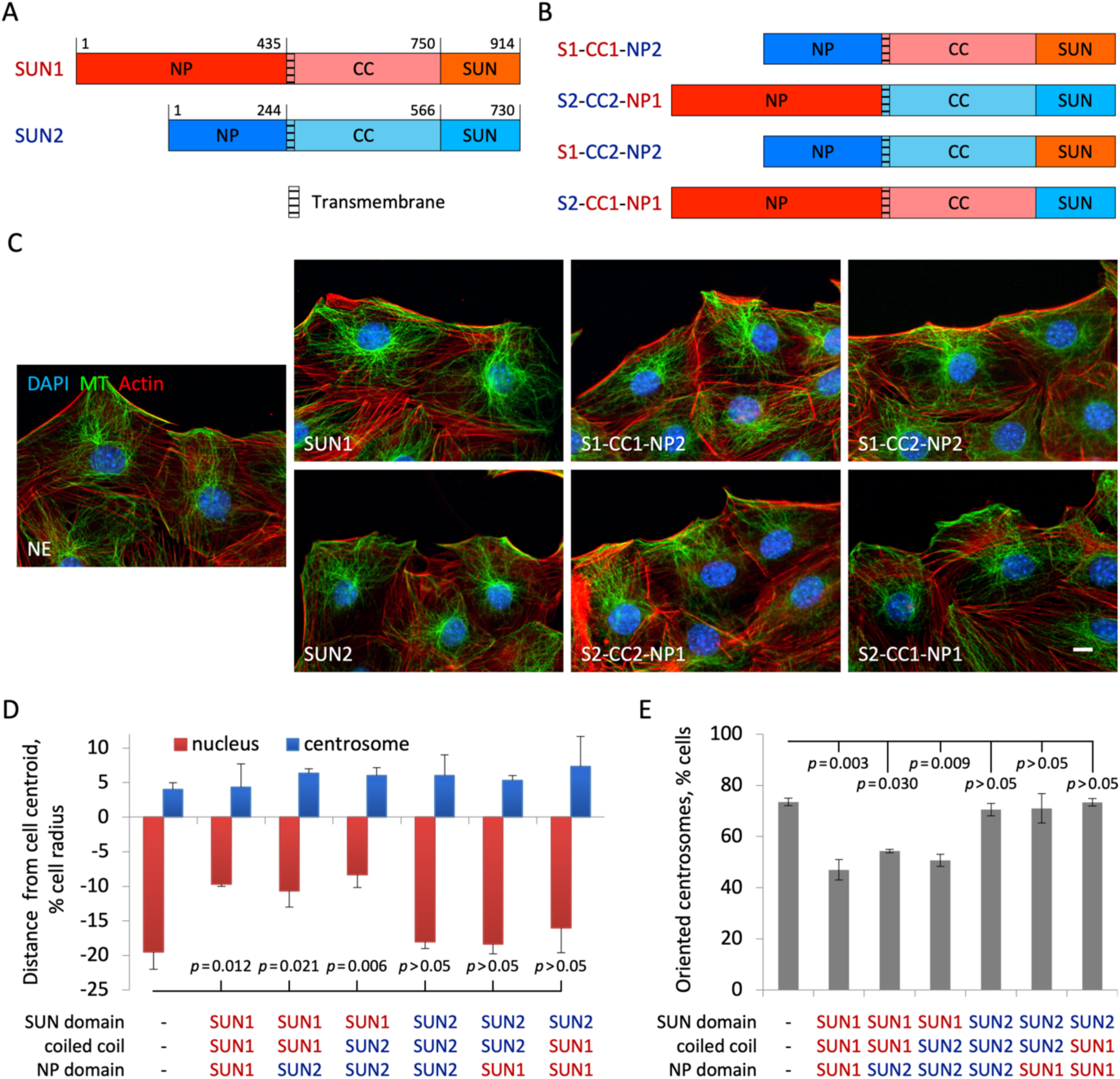
The SUN domain of SUN1 specifies its microtubule-dependent inhibitory effect on rearward nuclear movement. **A)** Domain structures of SUN1 and SUN2. NP, nucleoplasmic domain; CC, coiled-coil domain and SUN, SUN domain. **B)** Domain structures of chimeric SUN proteins. **C)** Representative images of LPA-stimulated non-expressing cells (NE) and cells expressing the indicated myc-tagged SUN proteins and chimeras. **D, E)** Nuclear and centrosome positions (D) and levels of oriented centrosomes (E) for cells as in panel C. Bar, 10 µm. *p*-values are by one-way ANOVA with a post-hoc Tukey test (*N* = 3, *n* ≥ 90).

We also examined the ability of these chimeric proteins to rescue actin-dependent nuclear movement in fibroblasts depleted of SUN2 (Fig. S8B, C), a crucial factor for nuclear movement during fibroblast migration [17,21]. Consistent with the notion that the SUN domains determine the roles of SUN proteins in nuclear movement, only the chimera with the SUN2 domain rescued the cell polarity defect in SUN2-depleted fibroblasts (Fig. 7). Together, these results show that the SUN domains of SUN proteins are not interchangeable and individually support either actin or microtubule functionality of nesprin-2G.

**Figure 7.**
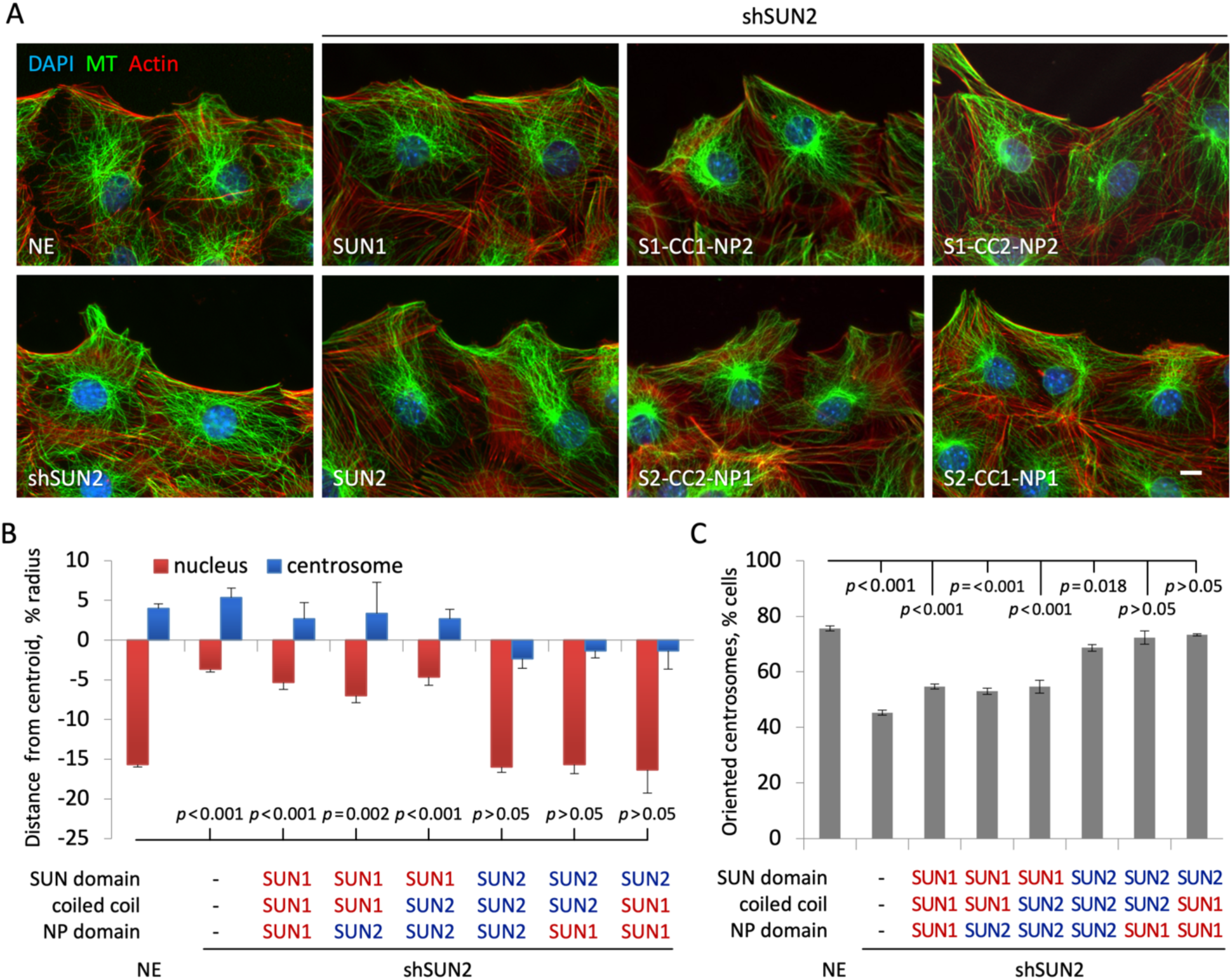
The SUN domain of SUN2 is required for actin-dependent rearward nuclear movement. **A)** Representative images of LPA-stimulated non-expressing cells (NE), cells expressing a SUN2-specific shRNA (shSUN2), and SUN2-knonkdown cells expressing the indicated myc-tagged SUN proteins and chimeras. **B, C)** Nuclear and centrosome positions (B) and levels of oriented centrosomes (C) for cells as in panel A. Bar, 10 µm. *p*-values are by one-way ANOVA with a post-hoc Tukey test (*N* = 3, *n* ≥ 90).

## Discussion

Despite early work suggesting that the two widely expressed SUN1 and SUN2 proteins can function redundantly, there is growing evidence for their specific functions [21,43]. One of the more striking examples is how SUN1 and SUN2 separately support microtubule- and actin-dependent positioning of the nucleus, respectively, in fibroblasts [21]. In this case, not only do the two SUN proteins support different cytoskeletal linkages to the nucleus, but when overexpressed, they transdominantly inhibit the linkage supported by the other SUN protein. This transdominant effect is observed not only in cases where SUN protein levels are manipulated experimentally but also underpins the defective cell polarization in physiologically and pathologically aged fibroblasts, where “naturally” elevated SUN1 expression inhibits actin- and SUN2-dependent positioning of the nucleus [23]. Elevated SUN1 is also observed in mouse models of aging, and reducing its expression ameliorates aged phenotypes and extends lifespan [22]. Whether the phenotypes in the mouse are caused by excessive microtubule interaction with the nucleus would be interesting to test.

Here, we demonstrate that the transdominant inhibition of SUN2 by SUN1 is due to its ability to act within the LINC complex. Using point mutants that disrupt the binding of SUN1 to lamin A and the covalent attachment of SUN1 to nesprins, we show that these interactions are essential for the transdominant activity. These interactions are key characteristics of SUN protein participation in the LINC complex, indicating that the transdominant effect is mediated primarily through the LINC complex. We provide further evidence that SUN1 specifically supports LINC complex connections to microtubules by showing that SUN1 expression enhances microtubule stability and that microtubule stability blocks actin-dependent nuclear movement in a SUN1-dependent manner. We identify the nesprin involved as nesprin-2 and highlight the involvement of its microtubule-interacting region. Lastly, we use chimeras of SUN1 and SUN2 to show that the specificity for microtubule and actin functionalities is determined by the SUN domain rather than the nucleoplasmic or coiled-coil domains. Thus, our data support a model in which the interaction of the SUN domain with nesprin-2 determines whether the nesprin will functionally interact with actin filaments or microtubules.

The dominant roles of the SUN domains in determining SUN protein’s functional interaction with the cytoskeleton are unexpected given that the SUN domains are highly conserved and the SUN1-KASH and SUN2-KASH structures closely match [32–34]. Considering that the SUN domains of SUN1 and SUN2 bind to the KASH peptide of nesprin-2 with virtually equal affinity [38], the specificity of nesprin-2G interaction with actin versus microtubules cannot be attributed to the tightness of binding. From a structural perspective, it is intriguing to consider how such similar structures in the lumen can differentially regulate the interactions of the cytoplasmic regions. Our results with the chimeras also provide insights into the functional importance of the oligomeric states of SUN proteins. SUN2 is trimeric [32]. SUN1 has been shown to form dimers and tetramers in native PAGE gels but appears trimeric in crystals [44,34]. Our finding that the coiled-coil regions of the SUN proteins are interchangeable suggests either: 1) SUN1 and SUN2 have similar oligomeric states; 2) the oligomeric states do not contribute to force transmission; or 3) additional mechanisms exist to oligomerize SUN1.

Both nesprin and SUN proteins have LINC-independent functions. For example, the SUN domain of SUN1 is dispensable for its roles in nuclear pore complex biogenesis and mRNA export [45,46], and the pro-apoptotic activity of nesprin-2 is not affected by the expression of GFP-KASH, which disrupts its interaction with SUN proteins [47,48]. However, for both the inhibition of nuclear movement and the regulation of microtubule stability, the SUN1 and nesprin-2 LINC complex is involved. The involvement of nesprin-2 indicates that cytoskeletal regulation by SUN1 might not stem from the LINC-independent repression of the SRF/Mkl1 pathway [49]. One possible mechanism involves the small GTPase RhoA. SUN1 has been shown to enhance RhoA activity in vascular smooth muscle cells [50]. It was reported to inhibit RhoA activity and focal adhesion assembly in HeLa cells, although conflicting evidence exists [43,51]. Given that SUN2 also affects RhoA and the cytoskeleton, further research is warranted to understand the roles of the LINC complexes in cytoskeletal regulation [43,50,39].

Although the microtubule cytoskeleton is responsible for the movement of the nucleus in a number of contexts [52], there is growing evidence that it can also play an inhibitory role, especially when the nucleus is moved by the actin cytoskeleton. In aged fibroblasts, the excessive interaction of microtubules with the nucleus presents a significant physical barrier to nuclear movement [23]. In the current study, we show that artificially stabilizing microtubules with Taxol significantly inhibits nuclear movement in a SUN1- and dynein-dependent manner, underscoring the crucial role of microtubules in these aging-related defects. Relatedly, recent research has identified excessive microtubule stabilization as a phenotypic marker of cellular senescence in epithelial cells and fibroblasts [53]. Given the diverse functions of microtubules and the growing evidence linking stable microtubules to aging, further investigation is needed to understand how microtubule dynamics and their interaction with the nucleus are misregulated during aging and how these contribute to the aging process.

## Methods and Materials

Sources of cell lines and reagents are listed in Table S1.

### Cell culture

NIH3T3 fibroblasts and HEK293T cells were maintained at 37°C with 5% CO_2_. NIH3T3 fibroblasts were cultured in DMEM with 10% bovine calf serum and penicillin-streptomycin. 293T cells were cultured in DMEM with 10% fetal bovine serum and penicillin-streptomycin. Cells were split with trypsin-EDTA when the cell confluency reached 80%. For serum starvation, confluent monolayers of cells on glass coverslips were washed three times with serum-free medium (DMEM with penicillin-streptomycin) and cultured in serum-free medium for 48 hours. Cells were tested for mycoplasma regularly. For drug treatments, HPI-4 was added at either 10 or 28 µM 1 h before LPA was added. Taxol (2 nM) treatment was 30 min and nocodazole (3.3 μM) treatment was 15 min. Remodelin (10 µM) treatment was 24 h.

### Plasmid construct

Plasmids of myc-SUN1, myc-lamin A, myc-progerin, myc-prelamin A, and miniN2G in the pMSCV vector and pSuper-shSUN1s were previously described [17]. Plasmids of GFP-tagged nesprin-2G fragment were previously described [21]. Myc-SUN1^C759A^, Myc-prelamin A^R527P^ and myc-progerin^R527P^ were made by overlap PCR using primers containing the mutation. Myc-SUN1ΔN was constructed from full-length myc-SUN1 with the 1-138 residues deleted. SUN1 and SUN2 chimeric plasmids were made by Gibson assembly. All the primer sequences are listed on Table S1.

### Retrovirus production and infection

2.5 µg pMSCV or pSuper plasmids were used to transfect HEK293T cells cultured on 35 mm dishes/wells, together with 2 µg pGag-pol and 1 µg pVSV-G plasmids using standard calcium phosphate transfection or Lipofectamine™ 3000 Transfection Reagent. Medium containing the viruses were collected at 48 and 56 h after transfection, pooled, aliquoted, and stored at −80 °C. 1 mL of virus and polybrene (4 µg/mL) were added to cells in 35-mm dishes/wells. Infected cells were selected using puromycin (1-10 µg/mL).

### Western blot

Cells were lyzed by adding RIPA buffer and kept on ice for 15 minutes. After adding sample buffer, the samples were boiled for 10 min and centrifuged at 13,000 rpm for 5 min. Proteins in the samples were separated by SDS-PAGE and transferred to nitrocellulose membranes. The membranes were blocked with TBS-T containing 5% non-fat milk for 1 h, incubated with primary at 4 °C overnight, washed, incubated with secondary antibodies for 1 h at room temperature, and washed. Images are acquired on a ChemiDoc MP Imaging system directly for dye-conjugated secondary antibodies, or after adding the ECL reagents for HRP-conjugated secondary antibodies. Images were processed and analyzed using Image J.

### Co-immunoprecipitation

Cells were lysed with milder lysis buffer (50 mM Tris, 150 mM NaCl, 1% NP-40, 0.25% sodium deoxycholate, pH 7.4) and the lysates were pre-incubated with IgG control and magnetic beads for 2-3 h at 4 °C. The supernatant was then co-incubated with control and primary antibodies and magnetic beads overnight at 4 °C. The pelleted beads were washed three times with the lysis buffer and resuspended in the original volume of lysis buffer. Samples were analyzed using Western blot.

### Cell migration (wound closure) assay

Monolayers of NIH 3T3 fibroblasts in DMEM with 2% Bovine Calf Serum were wounded and monitored on an EVOS FL LED Fluorescence Microscope. Images at different time points after wounding were analyzed using Image J.

### Cell polarization assay

Near confluent monolayers of NIH 3T3 fibroblasts were seeded on glass coverslips and serum-starved for two days. The starved monolayers were wound with a pipette tip and cells were allowed to recover for 30 mins. 10 µM LPA was added to stimulate nuclear movement and cell polarization for 2 h. Cells were then fixed with 4% PFA (room temperature, 10 mins). Cells were then immune-stained and images were acquired on a microscope (Nikon Ti2-E equipped with Qi-2 camera, objectives: 40x Plan-Fluro; 60x Plan-Apo) and analyzed with CellPlot [54].

### Immunofluorescence

Coverslips with fixed cells were washed three times with PBS and blocked in PBS with 0.3% Triton and 1% normal donkey serum for 1 h at room temperature. After incubation with the primary antibodies at 4 °C overnight, the coverslips were washed three times with PBS and incubated with secondary antibodies, phalloidin, and 4′,6-diamidino-2-phenylindole (DAPI) for 1 h. Stained coverslips were then washed and mounted on slides using Fluoromount-G medium.

### Statistics

Statistical analyses were performed using GraphPad. Unless noted otherwise, all values in the plots are mean ± SEM from *N* ≥ 3 independent experiments. For nuclear and centrosomal positions, at least 30 cells were measured for each condition in each experiment. Rates of oriented centrosomes are from at least 50 cells for each condition in each experiment. All Student’s *t*-tests are two-tailed and unpaired. Unless noted, unpaired one-way ANOVA tests are used, followed by either Tukey or Dunnett multiple comparisons test.

## Abbreviations

CC: coiled coil domain
HGPS: Hutchinson-Gilford Progeria Syndrome
HPI-4: Hedgehog pathway inhibitor-4
INM: inner nuclear membrane
KASH: Klarsicht, ANC-1, and Syne homology
LINC: linker of nucleoskeleton and cytoskeleton
LPA: lysophosphatidic acid
N2G: nesprin-2G
NAT10: N-acetyltransferase 10
NP: nucleoplasmic domain
ONM: outer nuclear membrane
SUN: Sad-1 and UNC-84

## Acknowledgments

This work was supported by funding from the Science and Technology Development Fund, Macau SAR (0077/2020/A2, 0099/2022/AFJ, and 0061/2022/A to W.C.), the University of Macau (MYRG2022-00251-FHS to W.C.), and NIH (R01 AG064944 and R35 GM136403 to G.G.G.).

