## Supplemental Figures and Table for "Elevated SUN1 promotes aging-related polarity defects through mechanical coupling microtubules to the nuclear lamina"

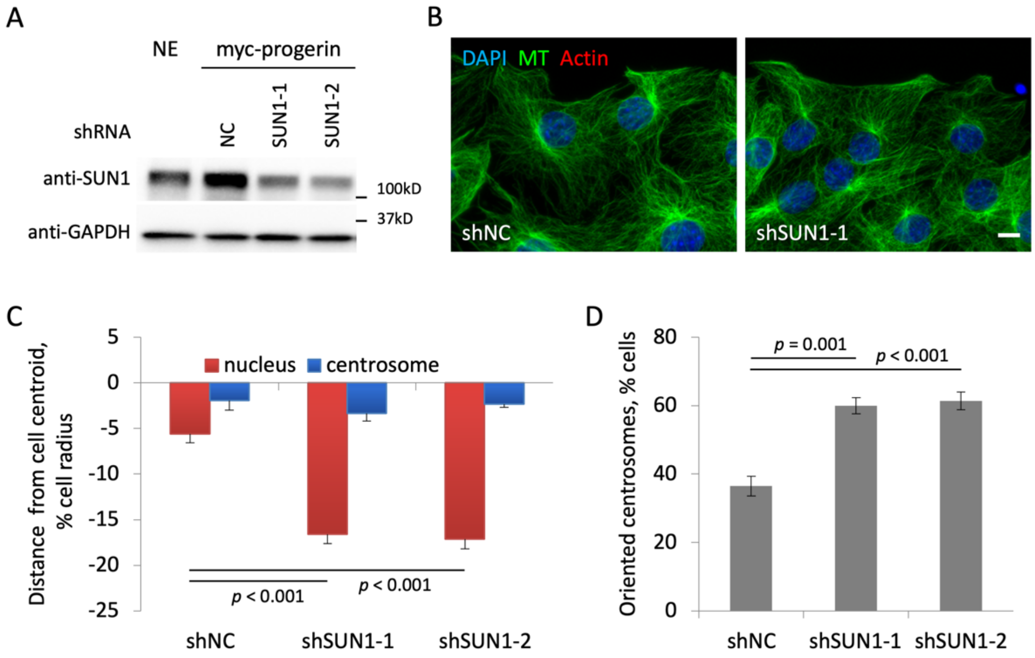


**Figure S1. Progerin expression inhibits nuclear movement through SUN1.**
**A)** Western blot of myc-progerin-expressing NIH3T3 fibroblasts showing the knockdown of SUN1 with shRNAs. NE: non-expressing cells; NC: noncoding shRNA. **B)** Representative images of serum-starved, LPA-stimulated monolayers of fibroblasts expressing the indicated shRNAs. **C, D)** Nuclear and centrosome positions normalized to cell size (C) and centrosome orientation (D) for fibroblasts expressing indicated shRNAs. Bar, 10 µm. *p*-values are by one-way ANOVA with a post-hoc Dunnett test ($N = 3$, $n \geq 90$).

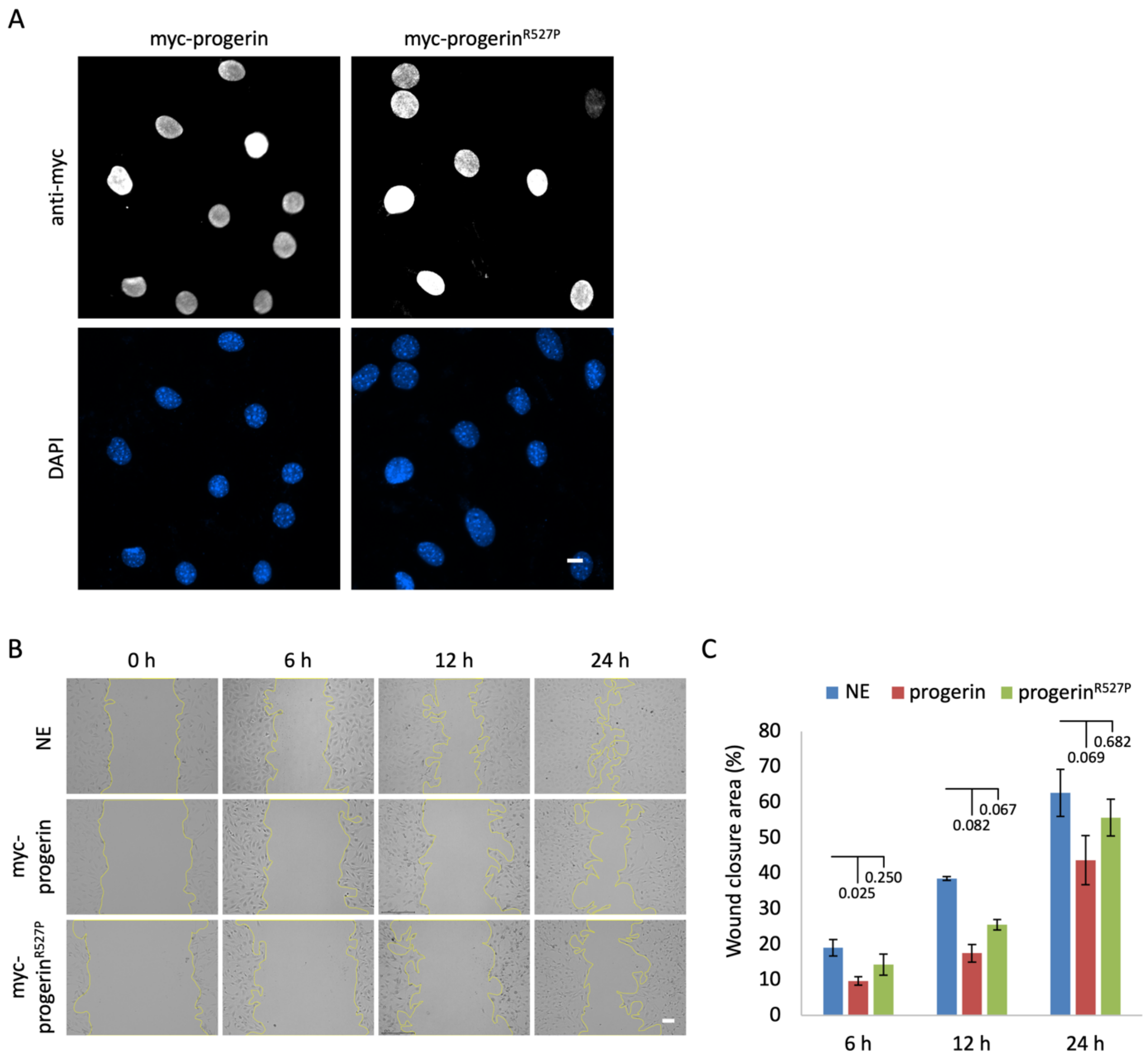


**Figure S2. Progerin$^{R527P}$ exhibits reduced association with SUN1 and improves cell migration when compared to progerin.**

**A)** Representative images showing the nuclear envelope localization of myc-progerin and myc-progerin$^{R527P}$ in NIH3T3 fibroblasts. Bar, 10 µm. **B, C)** Images (B) and quantification (C) of wound healing of non-expressing (NE) and cells expressing myc-progerin and myc-progerin$^{R527P}$. Bar in B, 100 µm. *p*-values are by one-way ANOVA with a post-hoc Dunnett test ($N = 3$).

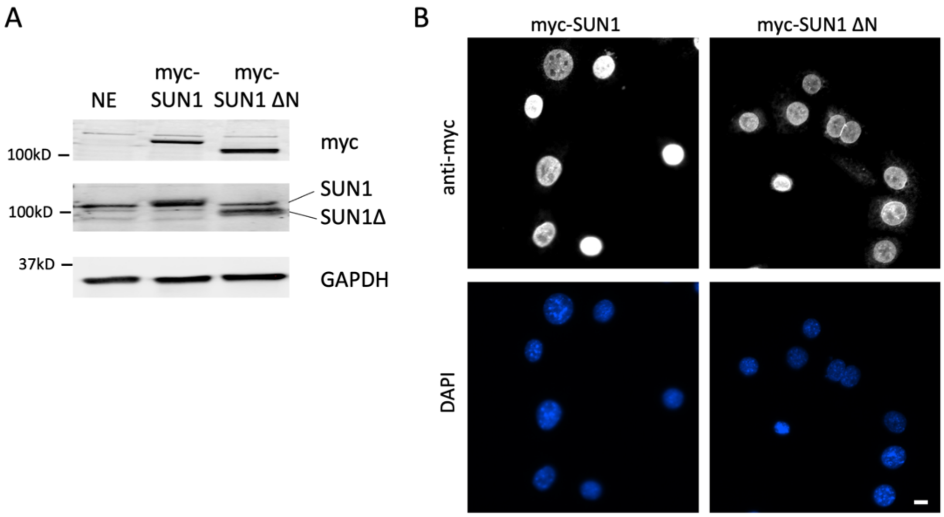


**Figure S3. SUN1ΔN localizes to the nuclear envelope.**
**A)** Western blot showing the expression of myc-SUN1 and myc-SUN1ΔN in NIH3T3 fibroblasts. NE: non-expressing. **B)** Representative images showing the nuclear envelope localization of myc-SUN1 and myc-SUN1ΔN in NIH3T3 fibroblasts. Bar, 10 µm.

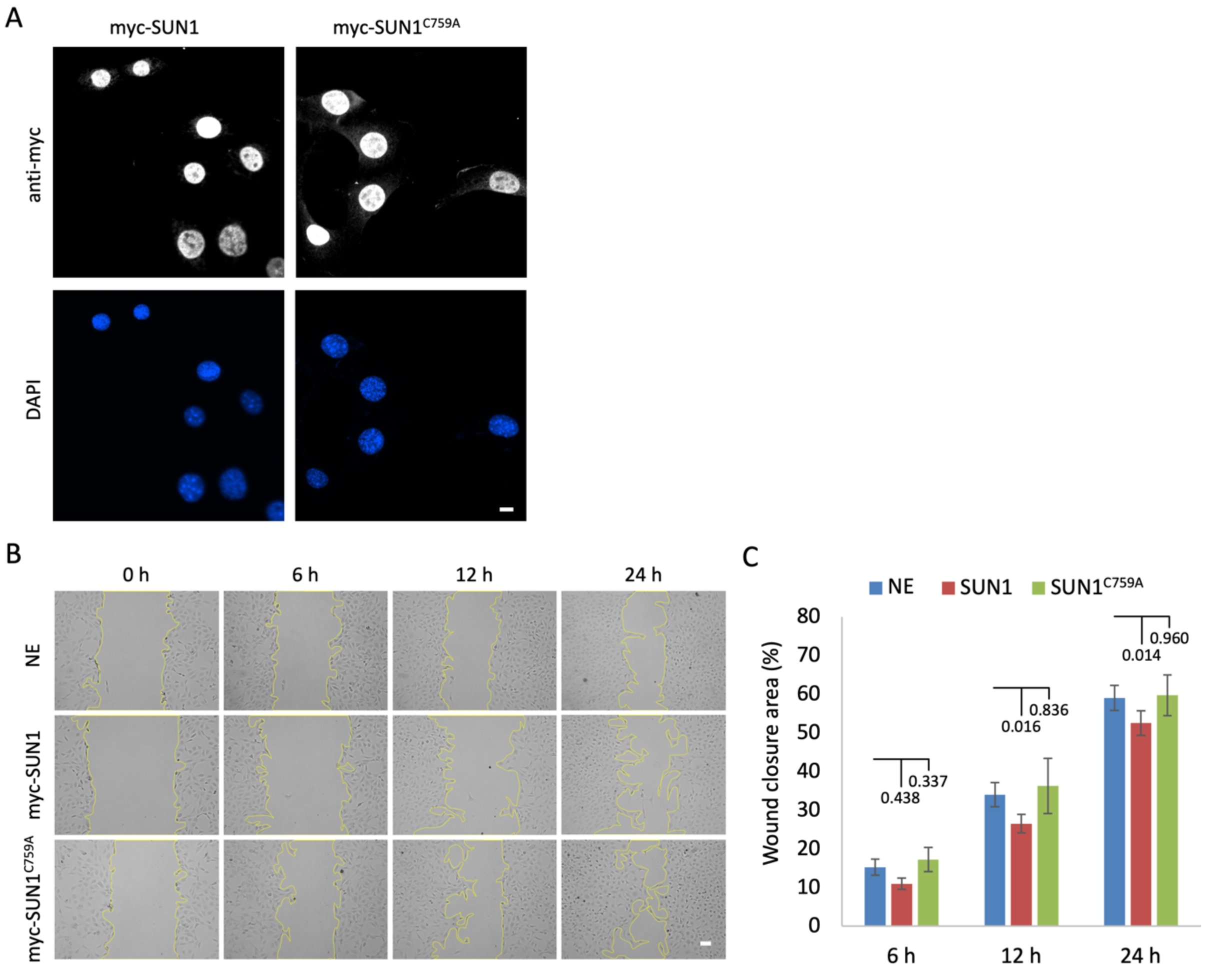


**Figure S4. SUN1$^{C759A}$ localizes to the nuclear envelope and does not interfere with cell migration.**

**A)** Representative images showing the nuclear envelope localization of myc-SUN1 and myc-SUN1$^{C759A}$.in NIH3T3 fibroblasts. Bar, 10 µm. **B, C)** Images (B) and quantification (C) of wound healing of non-expressing cells (NE) expressing myc-SUN1 and myc-SUN1$^{C759A}$. Bar in B, 100 µm. *p*-values are by one-way ANOVA with a post-hoc Dunnett test ($N = 3$).

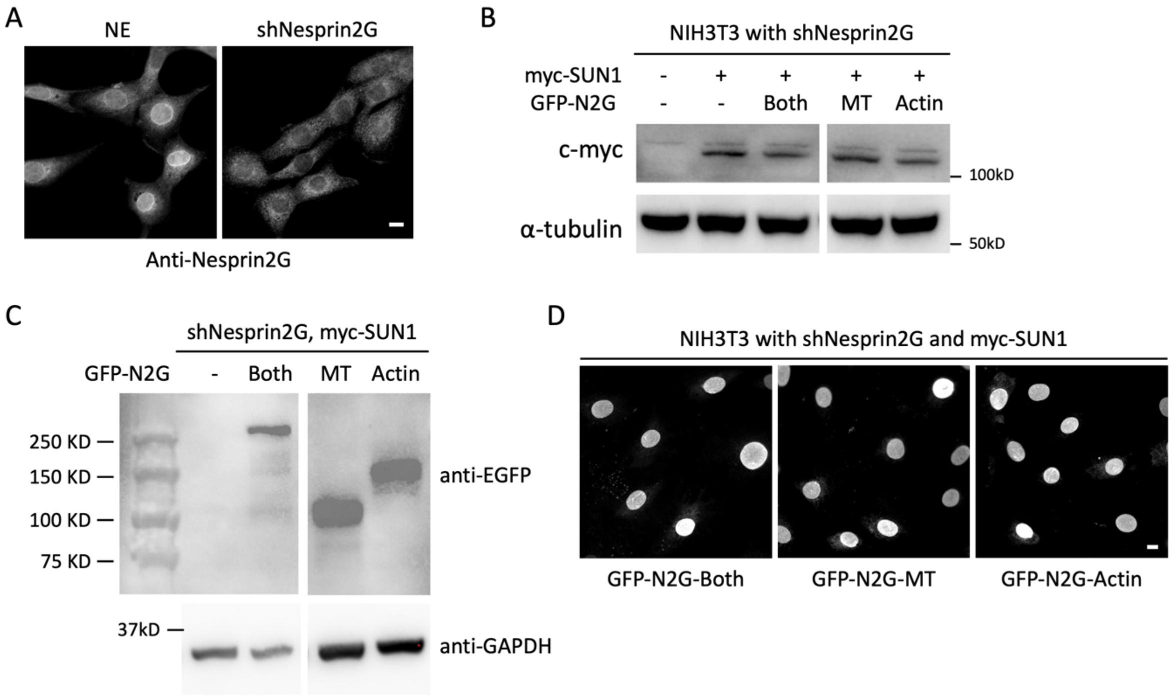


**Figure S5. shRNA depletion of nesprin-2G and expression of nesprin-2G fragments.**
**A)** Immunofluorescence images demonstrating the effective depletion of nesprin-2G, indicated by the absence of nuclear envelope staining. **B)** Western blot showing the expression of myc-SUN1. α-tubulin is a loading control. **C)** Western blot showing the expression of GFP-tagged nesprin-2G fragments in NIH3T3 fibroblasts expressing shNesprin-2G and myc-SUN1. **D)** All three nesprin-2G fragments exhibited nuclear envelope localization. Bars, 10 µm.

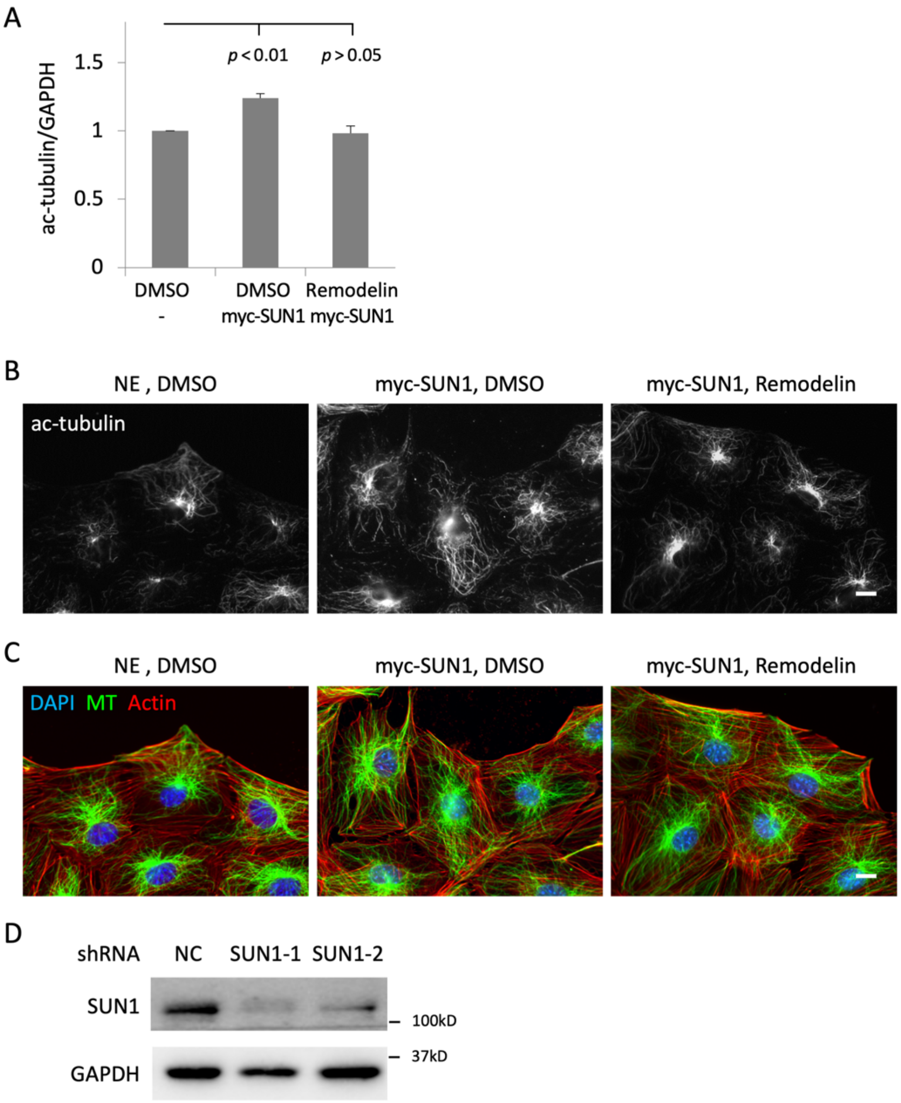


**Figure S6. Inhibition of stable microtubules restores rearward nuclear movement in a SUN1-dependent manner.**

**A)** Quantification of α-tubulin acetylation for samples in Figure S5A. *p*-values are by one-way ANOVA with a post-hoc Dunnett test ($N = 3$). **B)** α-tubulin acetylation in LPA-stimulated wildtype NIH3T3 fibroblasts (Ctrl) and myc-SUN1 expressing cells treated with either DMSO of remodelin (10 µM, 24 h). **C)** Cells in panel B were costained for α-tubulin, actin filaments, and DNA to visualize their polarity. **D)** Western blot showing the depletion of SUN1 in NIH3T3 fibroblasts. Bars, 10 µm.

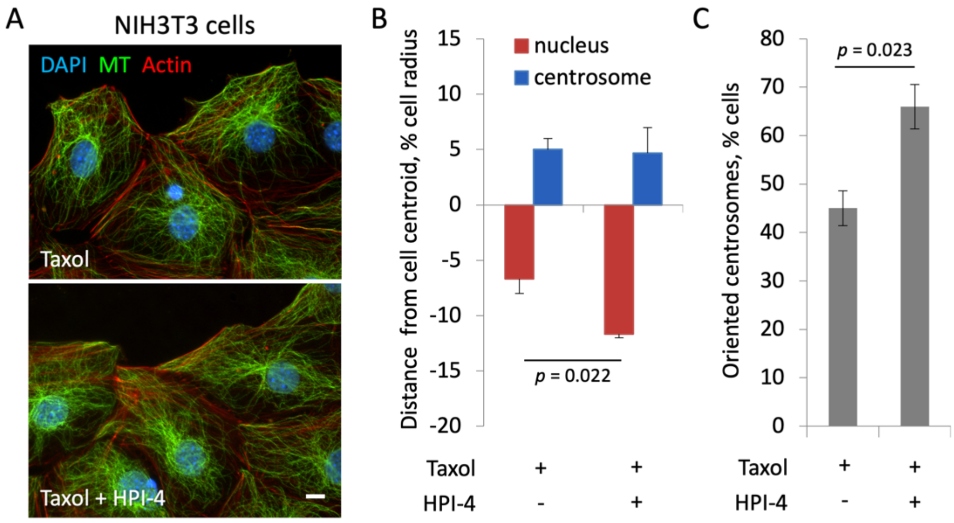


**Figure S7. HPI-4 reversed the suppression of cell polarity caused by Taxol.**
**A)** Representative images of wounded monolayers of LPA-stimulated NIH3T3 fibroblasts pretreated with Taxol (2 nM, 30 min) and either DMSO or HPI-4 (10 µM, 1 h). Bar, 10 µm. **B, C)** Nuclear and centrosome positions (B) and levels of oriented centrosomes (C) for cells as in panel A. *p*-values are by Student's t-test ($N = 3$, $n \geq 90$).

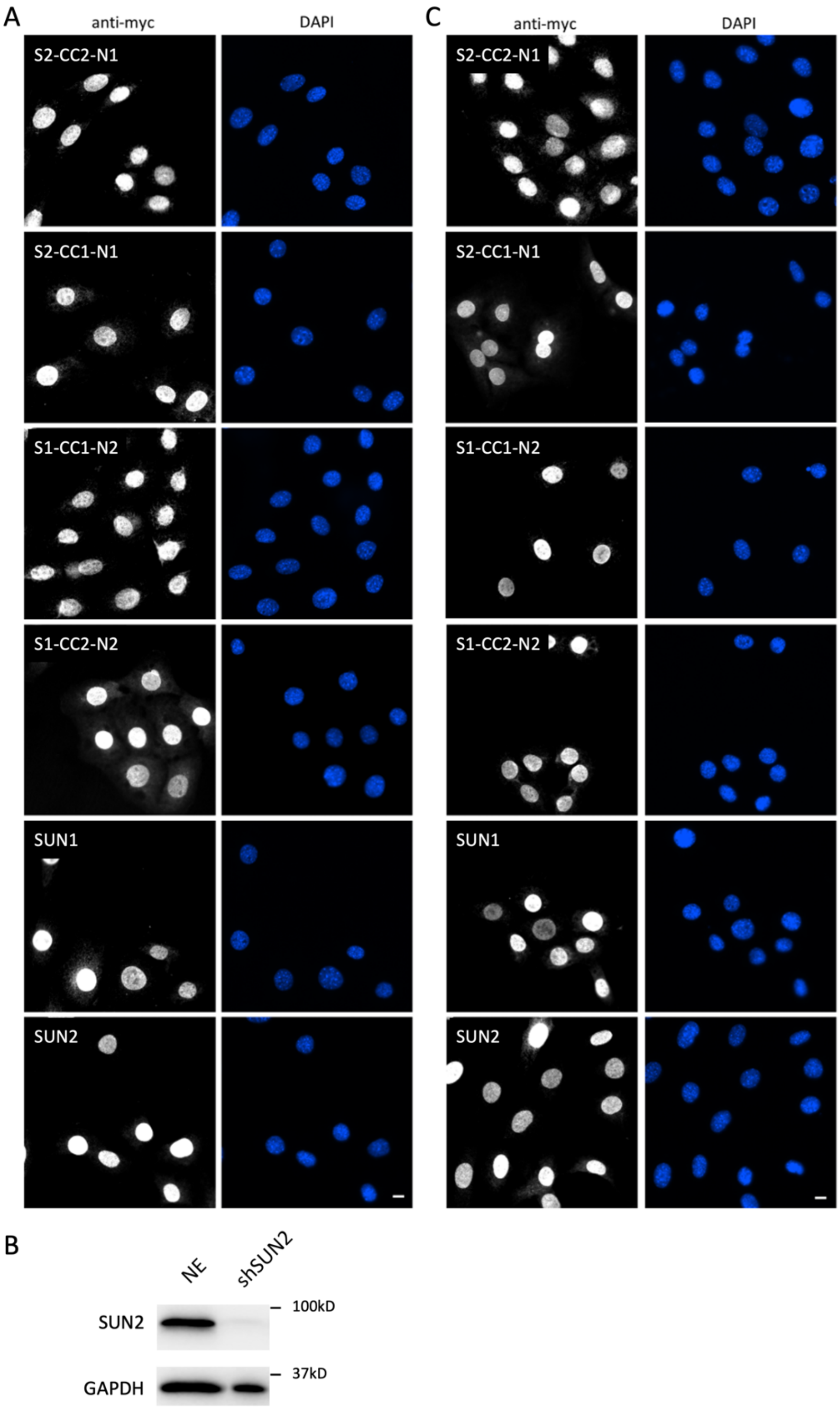


**Figure S8. Chimeric SUN proteins localize to the nuclear envelope.**

**A)** Representative images showing the nuclear envelope localization of myc-tagged chimeric and wildtype SUN proteins in NIH3T3 fibroblasts. **B)** Western blot showing the depletion of SUN2 by a shRNA in NIH3T3 fiblrblasts. **C)** Representative images showing the nuclear envelope localization of myc-tagged proteins in shSUN2-expressing NIH3T3 fibroblasts. Bars, 10 μm.

### Table S1. Reagents

| Primers | Sequence |
|---|---|
| PMSCV-Universal-Forward | GGACTTGAATGAAAGATCTGCTGG |
| PMSCV-Universal-Reverse | TCCCCTACCCGGTAGAATTC |
| LaminA R527P-Forward | AACAGCCTGCCTACGGCTCTC |
| LaminA R527P-Reverse | GAGAGCCGTAGGCAGGCTGTT |
| SUN1-ΔN-Forward | CGCGGATCCATGTCCCTGATTCGTGAGCAGA |
| SUN1-ΔN-Reverse | TCCCCTACCCGGTAGAATTCGTT |
| F-Forward | GAAGAGGACTTGAATGAA |
| F-Reverse | ACTGCCTTGGGAAAAG |
| SUN1-TM-Forward | TTCTCACTCCTACCAGTG |
| SUN1-TM-Reverse | CACTGGTAGGAGTGAGAA |
| SUN1-Domain-Forward | GGGATTTCAGGAATCACA |
| SUN1-Domain-Reverse | TGTGATTCCTGAAATCCC |
| SUN2-TM-Forward | TTCTCACTCCTACCAGTGTTAGGGCTGCAGACATTG |
| SUN2-TM-Reverse | GTGAGGTAGGAGTGAGAACAATGTCTGCAGCCCTAA |
| SUN2-Domain-Forward | GGGATTTCAGGAATCACAATAGTTGGGGTGACAGAG |
| SUN2-Domain-Reverse | TGTGATTCCTGAAATCCCCTCTGTCACCCCAACTAT |
| **Enzymes** | **Source and catalog number** |
| BamHI-HF | New England Biolabs, R3136T |
| EcoRI-HF | New England Biolabs, R3101T |
| NotI-HF | New England Biolabs, R3189L |
| BglI | New England Biolabs, R0143L |
| Q5® High-Fidelity2X Master Mix | New England Biolabs, M0492L |
| T4 DNA Ligase | New England Biolabs, M0202T |
| **Plasmid** | **Source** |
| pCMV-GagPol | This paper |
| pLP VSVG | This paper |
| pMSCV-puro-C-Myc-prelaminA | This paper |
| pMSCV-puro-C-Myc-progerin | This paper |
| pMSCV-puro-C-Myc-SUN1 | This paper |
| pMSCV-puro-C-Myc-SUN1$^{C759A}$ | This paper |
| pMSCV-puro-C-Myc-SUN2 | This paper |
| pMSCV-puro-EGFP-C4-miniN2G | This paper |
| pSuper-puro-SUN1, targeting sequence: CTGTATTTGACTCTCCA | This paper |
| pSuper-puro-SUN1, targeting sequence: | This paper |

| | |
|---|---|
| CCTTAAAGGAAATAAA | |
| pSuper-puro-SUN2, targeting sequence: GGGTCATTCTGCAGCCAGA | This paper |
| pMSCV-puro-C-Myc-prelaminA$^{R527P}$ | This paper |
| pMSCV-puro-C-Myc-progerin$^{R527P}$ | This paper |
| pMSCV-puro-C-Myc-SUN1ΔN | This paper |
| pMSCV-puro-C-Myc-S2-CC2-N1 | This paper |
| pMSCV-puro-C-Myc-S2-CC1-N1 | This paper |
| pMSCV-puro-C-Myc-S1-CC1-N2 | This paper |
| pMSCV-puro-C-Myc-S1-CC2-N2 | This paper |
| **Antibodies** | **Source and catalog number** |
| Anti-GAPDH antibody | BioLegend, 607902 |
| Anti-Myc/c-Myc Antibody (9E10) | Santa Cruz, sc-40 |
| Anti-SUN1 antibody (X12.11), monoclonal | Dr. Brian Bruke, A*STAR, Signapore |
| Anti-SUN1 antibody (X15.15), monoclonal | Dr. Brian Bruke, A*STAR, Signapore |
| Anti-SUN1 antibody (ab103021) | Abcam, ab103021 |
| Anti-UNC84A antibody | Invitrogen, PA5-113917 |
| Anti-SUN2 antibody (A-10) | Santa Cruz, sc-515330 |
| Anti-SUN2 antibody | Abcam, ab87036 |
| Anti-SUN2 antibody [EPR6557] | Abcam, ab124916 |
| Anti-lamin A/C antibody (E-1) | Santa Cruz, sc-376248 |
| Anti-acetylated α-tubulin antibody | Santa Cruz, sc-23950 |
| Anti-α-tubulin antibody (YL1/2) | European Collection of Animal Cell Cultures |
| Anti-α-tubulin antibody (DM1A) | Santa Cruz, sc-32293 |
| **Chemicals** | **Source and catalog number** |
| 4',6-Diamidino-2-Phenylindole, dihydrochloride | Invitrogen, D1306 |
| Alexa Fluor Plus 647 Phalloidin | Invitrogen, A30107 |
| Alexa Fluor Plus 555 Phalloidin | Invitrogen, A30106 |
| Lysophosphatidic acid (18:1) | Avanti Polar Lipids, 857130 |
| Ciliobrevin A (HPI-4) | MedChem Express, HY-100790 |
| Nocodazole | Sigma, M1404 |
| Taxol | Merck, PHL89806 |
| Remodelin | Sigma, SML1112 |
| HEPES | Aladdin, H109407 |
| Polybrene | Beyotime, C0351 |
| Puromycin (solution) | Invivo Gen, ant-pr-1 |
| Lipofectamine™ 3000 Transfection Reagent | Invitrogen, L3000075 |
| Glycine | Macklin, G800880 |

| | |
|---|---|
| Methanol, LC-MS Grade | Anaqua, MA-1296-4000 |
| Paraformaldehyde | Macklin, P804537 |
| Triton™ X-100 | Aladdin, T109026 |
| Tween 20 | Macklin, T6335 |
| LB Agar | Affymetrix, J75851.A1 |
| LB Broth | Affymetrix, J75852.30 |
| Fluoromount-G | Thermo Fisher Scientific, 00-4958-02 |
| RIPA buffer | Beyotime, P0013E |
| IgG from mouse serum | Sigma, I15381 |
| **Cell lines and tissue culture** | **Source and catalog number** |
| NIH3T3 cells | ATCC |
| HEK293T cells | Dr. Han-Ming Shen, Univ. Macau, Macau |
| DMEM, high glucose | GIBCO, 11965092 |
| Bovine calf serum | Cytiva, AH30203831 |
| FBS, Fetal Bovine Serum | GIBCO, 10270106 |
| Fatty lipid free Bovine Serum Album | Sigma, A6003 |
| Penicillin-Streptomycin | GIBCO, 15140163 |
| Trypsin-EDTA (0.25%), phenol red | Gibco, 25-200-056 |
| **Others** | **Source** |
| TOP10 Competent Cells | Dr. Qi Zhao, Univ. Macau, Macau |
| TIANprep Mini Plasmin Kit | TIANGEN, DP103-03 |
| SureBeads Protein G Magnetic beads | Bio-Rad, 1614023 |
